# A sensory map of the gastrointestinal tract

**DOI:** 10.1101/2025.03.07.641228

**Authors:** Reem Hasnah, Melanie Makhlouf, Matteo Zambon, Michela Arietti, Yolanda Graciela Kiesling Altún, Antonio Scialdone, Luis R. Saraiva, M. Maya Kaelberer

**Affiliations:** Research Branch, Sidra Medicine; Doha, Qatar; College of Health and Life Sciences, Hamad Bin Khalifa University; Doha, Qatar; Institute of Epigenetics and Stem Cells, Helmholtz Munich, 81377 Munich, Germany; Institute of Functional Epigenetics, Helmholtz Munich, 85764 Neuherberg, Germany; Institute of Computational Biology, Helmholtz Munich, 85764 Neuherberg, Germany; Trinity College of Arts & Sciences, Duke University, Durham, NC 27710; USA; Department of Medicine, Duke University, Durham, NC 27710; USA; Department of Physiology, University of Arizona, Tucson, AZ 85721; USA; Department of Comparative Medicine, Yale University School of Medicine, New Haven, CT, USA

## Abstract

The gut, like the skin, exhibits a spatially organized sensory system with distinct sensory maps. In the gut, regions are specialized for detecting and responding to diverse signals including immune, microbial, and nutrient or chemical stimuli. This study employed spatial transcriptomics to characterize the molecular landscape of gut sensory cells along the gastrointestinal tract, revealing regional differences in gene expression profiles associated with various sensory modalities. We found that genes related to hormone and neurotransmission expression varied along the gut length at different resolutions, suggesting a division of labor where hormones mediate broader effects and neurotransmission enables rabid, localized signaling. Immune-related genes also exhibit spatial patterns, with increased expression towards the colon, consistent with the higher microbial density in this region. Furthermore, nutrient sensing is spatially regulated, with differential expression of receptors for fats, sugars, proteins, vitamins, and minerals along the gut. This spatial map of gut sensory cell gene expression provides a foundation for deciphering the complex code underlying gut-brain communication.

## MAIN

The gastrointestinal tract exhibits significant regional diversity, with each segment playing specialized roles in digestion, absorption, and secretion. While this functional diversity is generally known, the spatial molecular variations of gut sensory cells is unknown. These gut sensory cells, while a minority of the total epithelial cell population, significantly contribute and influence diverse physiological processes such as hormone secretion, nutrient sensing, and gut-brain communication.^1–5^ Characterizing the unique molecular signatures the gut’s sensory properties across different intestinal regions is crucial for understanding how the gut is identifying and filtering signals that are ultimately communicated to the brain. This knowledge gap underscores the need for a comprehensive analysis of gut sensory cell spatial organization to decipher the nuanced interactions between these cells and their microenvironment.

Emerging evidence suggests distinct transcriptional profiles for gut sensory cells based on their anatomical location. For instance, the proximal small intestine is exposed to a different luminal environment from the colon^6^, indicating the need for distinct sensory modalities. This spatial segregation aligns with previous findings demonstrating varying abundance of different gut hormones along the proximal to distal length of the gastrointestinal tract.^7–9^ One crucial aspect of gut sensory function is the ability to rapidly and accurately communicate with both local and distant targets. This communication relies not only on the release of hormones but also on rapid signaling through synaptic transmission. These specialized cells that communicate rapidly via synapse are called neuropod cells.^3,5^ The spatial organization of the molecular components underlying synaptic communication in gut sensory cells is a key area of investigation in the role of a gut sense to drive food choice.^5,10^ Understanding how these cells integrate hormonal and synaptic signaling throughout the gastrointestinal tract will lay the foundation for deciphering the complex neural circuits that regulate gut function.

Two critical roles of gut sensory cells are to monitor the gut’s immunological landscape and sense ingested nutrients. The gut epithelium is at the interface between the host and the external environment, constantly exposed to a diverse array of potential immune challenges, including food antigens, commensal microbes, and pathogens. ^11,12^ Gut sensory cells contribute to both local immune regulation and systemic immune responses.^13,14^ The spatial distribution of immune-related molecules within gut sensory cells is investigated in this study. In addition,, the gut must continuously monitor the nutritional content of the food we eat. Gut sensory cells are equipped with a diverse repertoire of nutrient receptors, allowing them to detect and respond to a wide range of nutrients.^15,16^ The spatial expression of these nutrient receptors throughout the gut likely reflects the region-specific functions absorption, metabolism, and food-related behaviors. Understanding how gut sensory cells integrate information from the immune system and about nutrient availability with other sensory inputs is essential for understanding the dependencies between diet, gut physiology, and overall health.

In this study, we employed spatial transcriptomics to create a detailed molecular map of gut sensory cells throughout the small intestine and colon. By examining the spatial expression patterns of genes involved in endocrine and synaptic signaling, immune function, and nutrient sensing, we aimed to uncover the regional specializations of these crucial cells and provide a comprehensive resource for future investigations into gut sensory biology.

## Results

### Profile of spatial transcriptomics of the small intestine

To optimize digestion and nutrient absorption, the gut demonstrates physiological differences along its length, resulting from variations in tissue architecture and cellular composition. To investigate the spatial gene expression patterns throughout the small intestine, the first 12-cm were divided into equal 1 cm segments (n=5 per segment) for RNA sequencing (Figure 1A). We found 7,179 genes with a differential expression pattern across these intestinal segments (see Methods). A hierarchical clustering algorithm identified eight distinct gene expression patterns characterized by four primary shapes: monotonically decreasing (\), inverted U-shaped (∩), U-shaped (∪), and monotonically increasing (/) (Figure 1B). More stringent criteria yielded similar pattern shapes but with fewer differentially expressed genes (Supp. Fig. 1A, B). Gene counts across patterns ranged from 331 genes in Pattern 3 to 2,180 genes in Pattern 1 (median of 744 genes per cluster).

**Figure 1.**
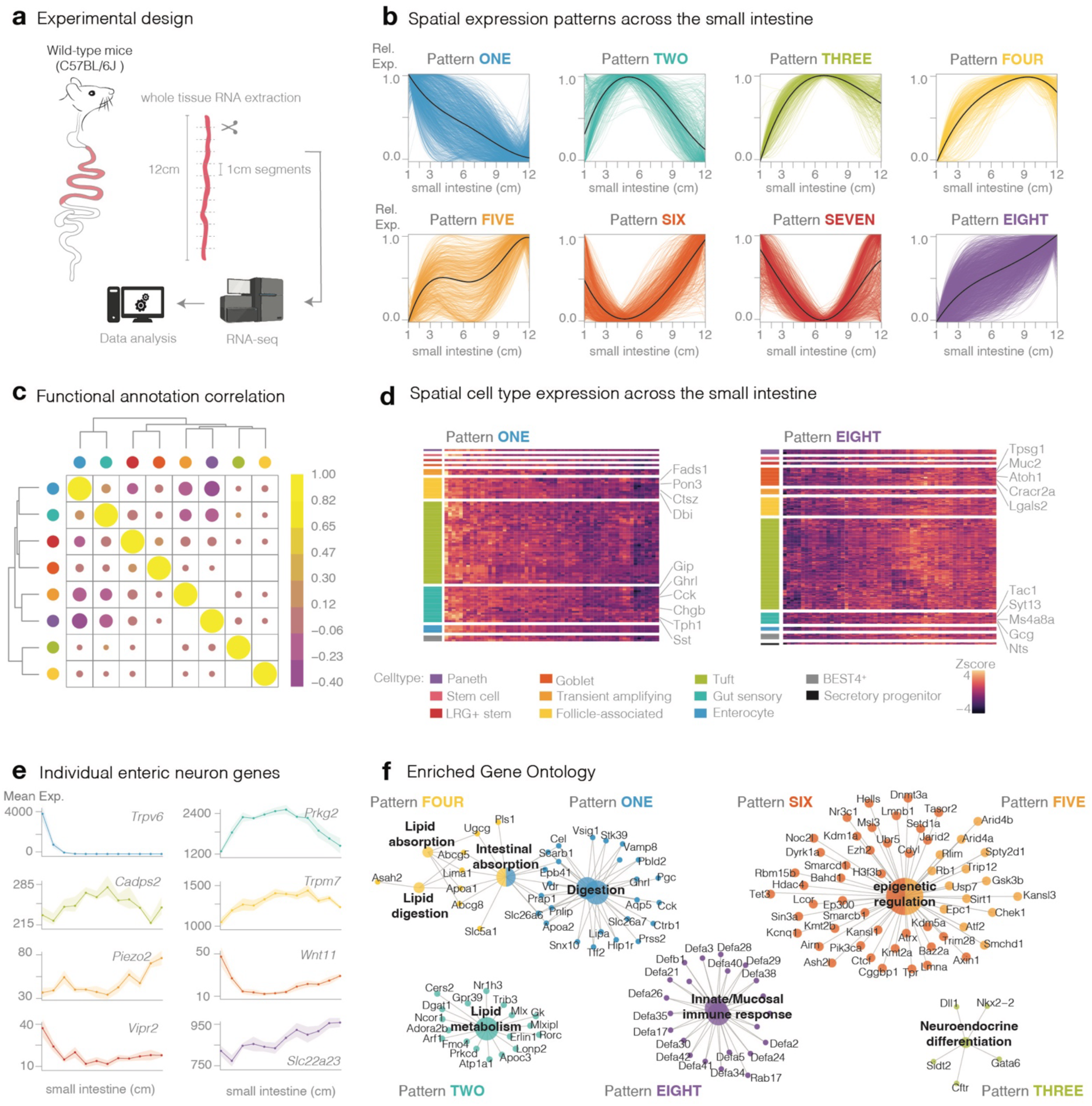
A transcriptional profile of small intestinal segments. (A) Schematic of the experimental design: The intestine was divided into twelve equal 1 cm segments from each sample (n=5), and RNA sequencing was performed on each segment. (B) Distinct patterns of differential gene expression along the small intestinal axis. Relative expression of individual patterned genes in color, with black line denoting average expression. (C) Spearman’s correlation of functional similarity between gene annotations within each pattern. (D) Intestinal cell lineage marker enrichment: Z-score heatmap illustrating key intestinal cell lineage markers enrichment within spatial Pattern 1 and 8. (E) Line plot of mean expression of enteric neuron marker genes across intestinal segments, mean with shaded SEM. (F) Gene Ontology (GO) enrichment map plot of selected gene ontologies.

Functional correlation shows that Patterns 1 and 8 were the most distinct (Figure 1C), highlighting their distinct regional specialization. Focusing on the monotonically decreasing Pattern 1 and the monotonically increasing Pattern 8, we examined the spatial expression patterns of intestinal cell types (Figure 1D). Based on previous literature, ^17–19^ all eleven cell types were represented, however, the Tuft cell type had the highest proportion of spatially regulated genes with a striking spatial segregation, where distinct populations favored either the proximal (Pattern 1) or distal (Pattern 8) small intestine. These findings align with recent evidence showing distinct Tuft cell populations that are immune-reactive.^20–22^ Additionally, gut sensory cell markers were the second most represented cell type in both spatial patterns, indicating that these cells are spatially specific. In this context, gut sensory cell markers *Gip*, *Ghrl*, *Tph1*, and *Sst* were enriched in Pattern 1, while *Tac1*, *Gcg*, and *Nts* were enriched in Pattern 8. This spatial heterogeneity likely reflects the region-specific functional demands of the small intestine; however, these are whole intestinal segment samples and individual genes could be expressed in non-epithelial cell populations. Therefore, additional isolation methods will be needed to better distinguish epithelial cell type distribution.

Given the abundance of enteric neurons in whole tissue samples, we also examined the specific spatial expression patterns based on previously published cluster analysis of all enteric neuron classes (ENC)^23^ (Supp. Fig. 1C). In contrast to epithelial cell types, no one enteric neuron class was associated with a single specific spatial pattern, suggesting a more consistent distribution of enteric neuron type throughout the small intestine. However, there were differences in the sub-types of enteric neurons based on single gene expression that was found to be spatially regulated genes (Figure 1D). For example, *Trpv6* (associated with the interneuron class, ENC10) was enriched in Pattern 1, while *Prkg2* (also ENC10) was enriched in Pattern 2. Indicating some spatial specificity within an enteric neuron class. These data suggest that a combinatorial understanding of cell type, location, and function is essential for a comprehensive understanding of intestinal physiology.

Kyoto Encyclopedia of Genes and Genomes (KEGG) pathway and Gene Ontology (GO) enrichment analyses were performed to explore the functional implications of these spatial gene expression patterns (Supp. Fig. 1D-F). KEGG pathways were categorized into Organismal Systems, Environmental Information Processing, and Metabolism. Metabolic pathways were strongly linked to Pattern 1, suggesting that the proximal small intestine plays a dominant role in nutrient breakdown. In contrast, Pattern 8 was enriched in pathways related to Environmental Information Processing, particularly microbial identification and response, indicating a greater emphasis on microbe-host interactions in the distal small intestine. GO enrichment analysis further refined these functional assignments. Patterns 1, 2, and 4 demonstrated a progression of enrichment for digestion, metabolism, lipid metabolism, then absorption moving from the proximal to distal small intestine. Patterns 5, 6, and 8 were associated with environmental cues and epigenetic regulation, likely reflecting responses to those cues in the distal small intestine. Pattern 3, peaking midway along the intestine, was linked with neuroendocrine differentiation, potentially indicating a role in nutrient sensing and signaling (Figure 1F). Taken together, these data demonstrate that spatially distinct cell populations adapt to perform specialized roles in cellular signaling, immune interactions, and nutrient sensing along the length of the small intestine.

### The molecular barcodes of Neurod1-labeled gut sensory cells

*Neurod1,* a transcription factor essential for establishing and promoting the differentiation of the neuroendocrine cell lineage,^24^ is expressed in all gut sensory cells and absent in other intestinal epithelial cell types.^25^ To investigate the molecular profile of these sensory cells across the gastrointestinal tract, we divided the Neurod1Cre_tdTomato mouse gut into three regions: the first half of the small intestine (1^st^ half), the second half of the small intestine (2^nd^ half), and the colon. Epithelial cells from each region were then sorted based on tdTomato fluorescence into Neurod1-labeled (sensory) and negative (non-sensory) populations for RNA-sequencing (Figure 2A). As expected, *Neurod1* expression was enriched in the Neurod1-labeled population and cell number decreased with length consistent with a decrease in overall surface area (Supp. Fig. 2A,B). However, number of cells and normalized counts had a positive correlation (Spearman correlation, R² = 0.8, p=2e-05), indicating a decrease in expression level from the proximal small intestine to the colon (Supp. Fig. 2C). In addition, Neurod1-labeled cells encompassed populations across all differentiation stages ^26,27^ (Supp. Fig. 2D). Principal component analysis shows that the main group difference is between Neurod1-labeled versus negative cells, followed by all small intestine versus colon epithelial cells (Figure 2B).

**Figure 2.**
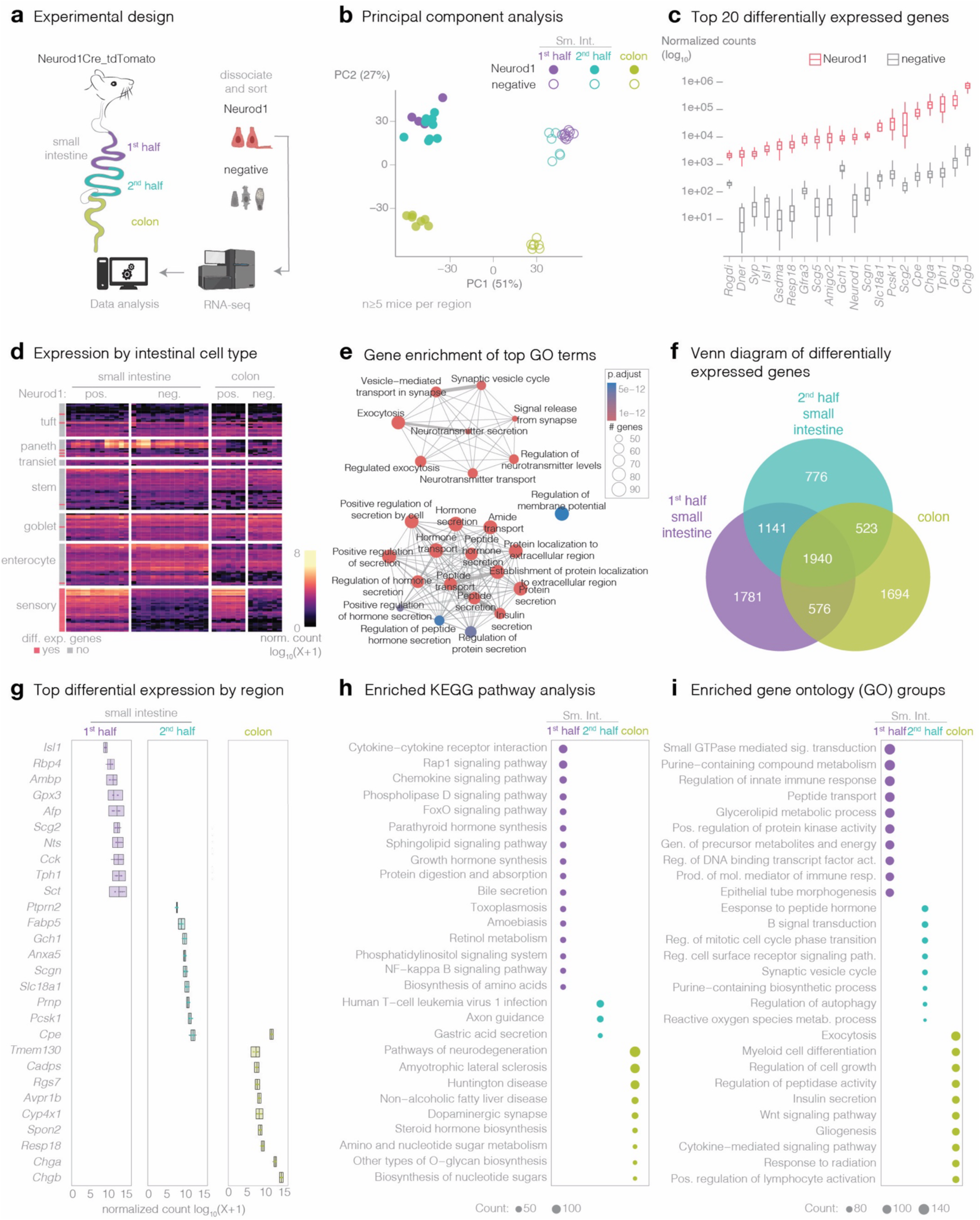
A transcriptional map of Neurod1-labeled cells of the small intestine and colon. (A) Experimental design of samples from Neurod1Cre_tdTomato mice taken from the length of the gastrointestinal tract, sorted and RNA-sequenced. (B) Principal Component Analysis (PCA) with visualization of the top 1000 most expressed genes across all samples (n= 4-8 mice per region). (C) Box plot of normalized counts for the top 20 differentially expressed genes between Neurod1-labeled and - negative cells. (D) Expression heatmap of marker genes for intestinal cell types in Neurod1-labeled and - negative cells. Each row represents cell type gene. Color intensity indicates relative expression. (F) Venn diagram of Neurod1-labeled differentially expressed genes from different locations. Numbers indicate uniquely expressed genes in segment. (G) Box plot of top differentially expressed genes between Neurod1-labeled versus negative cells by regions. Normalized expression counts ±SEM. (H) Dot plot of unique KEGG pathways enriched in Neurod1-labeled versus negative cells by region (small intestine: 1^st^ half:1781 genes, 2^nd^ half:776 genes, and colon:1694 genes). Each dot represents a unique KEGG pathway, with color indicating the specific region and size reflecting the number of genes enriched in the pathway. (I) Dot plot of unique gene ontology (GO) terms associated with differential expression in Neurod1-labeled versus negative cells by region (small intestine:1^st^ half:1781 genes, 2^nd^ half:776 genes, and colon:1694 genes). Each dot represents a unique GO term, with size reflecting the number of associated genes and color indicating the corresponding region.

Differential gene expression analysis of the combined Neurod1-labeled versus negative cell populations across the small intestine and colon identified three main functional groups among the top 20 differentially expressed genes: 1) transcription factors for sensory cell development (*Dner*, *Isl1*, *Neurod1*); 2) genes related to neuropeptide storage and secretion (*Scgn*, *Pcsk1*, *Cpe*, *Chga*, *Chgb*); and 3) genes associated with neuronal development and function (*Syp*, *Gfra3*, and *Slc18a1*) (Figure 2C). Many known gut sensory cell marker genes were enriched in Neurod1-labeled cells, while markers for other epithelial cell types were enriched in the negative population (Figure 2D). Further analysis, including a heatmap of specific marker gene enrichment, confirmed that the Neurod1-labeled population encompasses all identified gut sensory cell subtypes (Supp. Fig. 2E).

KEGG pathway and GO enrichment analyses were performed to evaluate the functional roles of Neurod1-labeled versus negative cells. The KEGG pathway analysis identified enrichment for the neuroactive ligand-receptor interactions pathway (mmu04080) (Supp. Fig. 2F), reflecting the importance of hormones, neuropeptides, neurotransmitters, and their receptors in gut sensory cell communication. Enrichment was also observed in the cell adhesion molecules pathway (mmu04514). Within this pathway, we identified genes involved in neuronal development (*Nfasc, Cntnap2*), axon guidance (*Ncam1*, *Ntng1*, *Ptprs*), myelin formation (*Mag*), and synapse formation (*Nlgn1*). Consistent with these findings, GO analysis revealed enrichment for gene ontologies related to hormone secretion, neurotransmitter signaling, and regulation of membrane potential, which are essential characteristics of electrically active cells (Figure 2E).

### Regional specificity of Neurod1-labeled cells

In all Neurod1-labeled cells, 1,940 genes were differentially expressed in all regions, 1,781 were unique to the first half of the small intestine, 776 were unique to the second half of the small intestine, and 1,694 were unique to the colon (Figure 2F). These data indicate a core set of sensory cell transcripts, with regional specializations, potentially with the second half of the small intestine representing a transitional zone between proximal and distal regions. To assess regional functional differences, we performed differential gene expression of Neurd1-labeled versus negative cells, KEGG pathway, and GO analyses (Figure 2G-I). Neurod1-labeled cells in the first half of the small intestine were enriched for genes associated with proximal gut hormones like cholecystokinin (Figure 2G), metabolic and absorptive pathways (mmu04976, mmu04974, mmu04071, mmu01230, and mmu04971) (Figure 2H), and neuronal functions, such as neurotransmitter transport and learning/memory (Figure 2I). In contrast, colon Neurod1-labeled cells were enriched in KEGG pathways associated with neurodegenerative diseases, including Parkinson’s disease (mmu05012) and Alzheimer’s disease (mmu05010), and GO terms related to regulating developmental growth, maturation, and social behavior (Figure 2H-I). These findings position the colon as a potential target for age-related biomarker discovery and interventions.

### The molecular barcodes of Cck-positive gut sensory cells

Cholecystokinin (Cck)-expressing gut sensory cells are a crucial subset located in the duodenum and jejunum.^8,28^ Their proximal location in the small intestine allows for rapid assesment of nutrient content to influence food choice^3,4^. To investigate the reginal specificity of these cells, we collected Cck-labeled cells from CckCre_tdTomato mice. The first two-thirds of the small intestine was divided into 3-cm increments classified as: proximal duodenum, distal duodenum, proximal jejunum, and distal jejunum (Figure 3A). Principal component analysis (PCA) shows a clear separation of Cck-labeled versus negative cells along PC1 (Figure 3B). However, within the Cck-labeled populations, PC2 reflects a regional progression (Figure 3C). This separation along PC2 was associated with genes involved in diverse cellular and metabolic processes, suggesting distinct physiological functions. For example, among the top 50 genes along PC2, genes such as *Alpi*, *Akp3*, *Sis*, and *Gstm3* are associated with metabolic and detoxification pathways, while *Traf6*, *Il17re*, and *Icosl* suggest immune regulatory involvement, and *Vim*, *Cntrl*, and *Nup210* are involved in cytoskeletal organization and cellular structure. Unlike *Neurod1*, there was no correlation between number of cells sorted and *Cck* normalized counts (Supp. Fig. 3A-B), indicating that *Cck* expression may be more consistend throughout the differnt small intestine regions.

**Figure 3.**
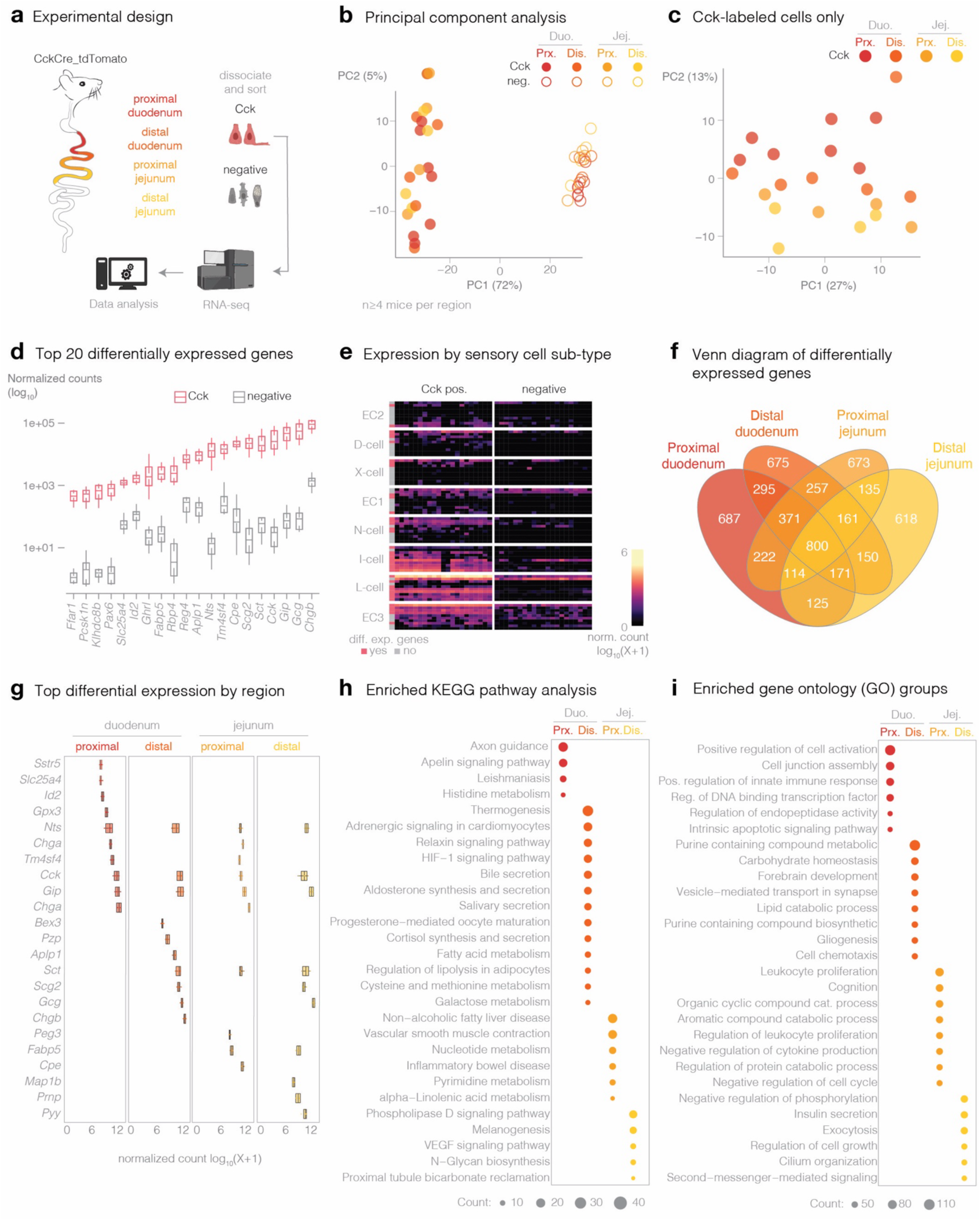
A transcriptional map of Cck-labeled cells of the first two-thirds of the small intestine (duodenum and jejunum). (A) Experimental design of samples from CckCre_tdTomato mice taken in 3-cm increments from P30-P40 mice of the first two-thirds of the small intestine, sorted and RNA-sequenced. (B) Principal Component Analysis (PCA) with visualization of the top 1000 most expressed genes in Cck-labeled and negative samples (n= 4-6 mice per region). (C) PCA of top 1000 most expressed genes across all Cck-labeled samples. (D) Box plot of normalized counts for the top 20 differentially expressed genes between Cck-labeled and -negative cells. (E) Expression heatmap of sensory cell subtype markers in Cck-labeled and -negative cells. Each row is a sensory cell subtype. Color indicates relative expression level. (F) Venn diagram of differentially expressed genes between Cck-labeled and negative cells by region. (G) Box plot of top differentially expressed genes between Cck-labeled versus negative cells by region. (H) Dot plot of top enrichment of KEGG pathways for differentially expressed genes by region (duodenum: proximal-687 genes, distal-675 genes; and jejunum: proximal-673 genes, distal-618 genes). Color indicates region and size the number of genes enriched in the pathway. (I) Dot plot of top Gene Ontology terms for differentially expressed genes by region (duodenum: proximal-687 genes, distal-675 genes; and jejunum: proximal-673 genes, distal-618genes). Color indicates region and size the number of genes.

Differential gene expression analysis between Cck-labeled and negative cells identified three main groups: 1) classic gut hormones (*Cck, Ghr1, Nts, Gip, Gcg*), suggesting potential co-secretion of multiple hormones^29–31^; 2) fatty acid sensing genes (Ffar1, Fabp5), highlighting the importance of these cells in lipid sensing; and 3) genes involved in differentiation and maturation (*Pax6, Tm4sf4, Reg4*), suggesting ongoing developmental specification within these cells (Figure 3D). A heatmap of genes associated with transcriptional stages confirms that Cck-labeled cells span a range of developmental stages (Supp. Fig. 3C). These findings challenge the "one-cell, one-hormone" dogma and underscore the need for a more comprehensive transcriptional and functional classification of Cck-labeled cell subtypes (Figure 3E).

KEGG pathway analysis revealed enrichment in multiple synapse-associated pathways, including serotonergic, dopaminergic, cholinergic, GABAergic, and glutamatergic synapses (Supp. Fig. 3D). This broad enrichment highlights a key role for Cck-labeled cells in rapid communication with neurons, also known as neuropod cells,^3,5^ via diverse signaling mechanisms.

### Regional specificity of Cck-positive cells

A Venn diagram of up-regulated regionally differentially expressed genes shows extensive overlap as well as discrete gene groups by region (Figure 3F). The top 10 differentially expressed genes in each region show relatively subtle differences between closely adjacent 3-cm segments (Figure 3G). Certain gut hormones (*Cck, Gip, Nts, and Sct*) were consistently expressed across regions, while others (*Gcg, Pyy*) showed more distal enrichment, suggesting a continuum of Cck-labeled cell expression profiles along the proximal-distal axis rather than discrete anatomical subdivisions. KEGG pathway and GO analyses were conducted to assess regional functional specificity (Figure 3H-I, Supp. Fig. 3E-F), using differentially expressed genes identified in each region: 2,785 (proximal duodenum), 2,880 (distal duodenum), 2,733 (proximal jejunum), and 2,274 (distal jejunum). KEGG pathway analysis revealed that Cck-labeled cells across all four segments are primarily oriented toward metabolic functions (Fig3 H-I). When the analysis was performed using differentially expressed genes from the entire duodenum (4,153 genes) and jejunum (3,391 genes), specific subcategories related to digestion and lipid metabolism (mmu00565, mmu04071) were identified in the duodenum (Supp. Fig. 3E-F).

### Spatial expression of hormone and neurotransmitter communication

Gut sensory cells, also known as neuropod cells, exhibit a dual functionality, acting as both endocrine cells releasing hormones and as specialized sensory cells engaging in synaptic transmission (Figure 4A). Volcano plots of both Cck- and Neurod1-labeled cells demonstrate a strong enrichment of hormone and neurotransmission genes in gut sensory cells compared to other epithelial cells (Figure 4B). To investigate regional transcriptomic differences in these cells, we identified spatially regulated genes associated with gut hormone production, neurotransmission, and synaptic structure. We performed differential expression analyses on samples collected from different locations within the Cck-labeled and Neurod1-labeled cell populations. Genes were then grouped based on their spatial expression patterns, classified by the sign of the log-fold change when comparing adjacent spatial sections (see *Methods*).

**Figure 4.**
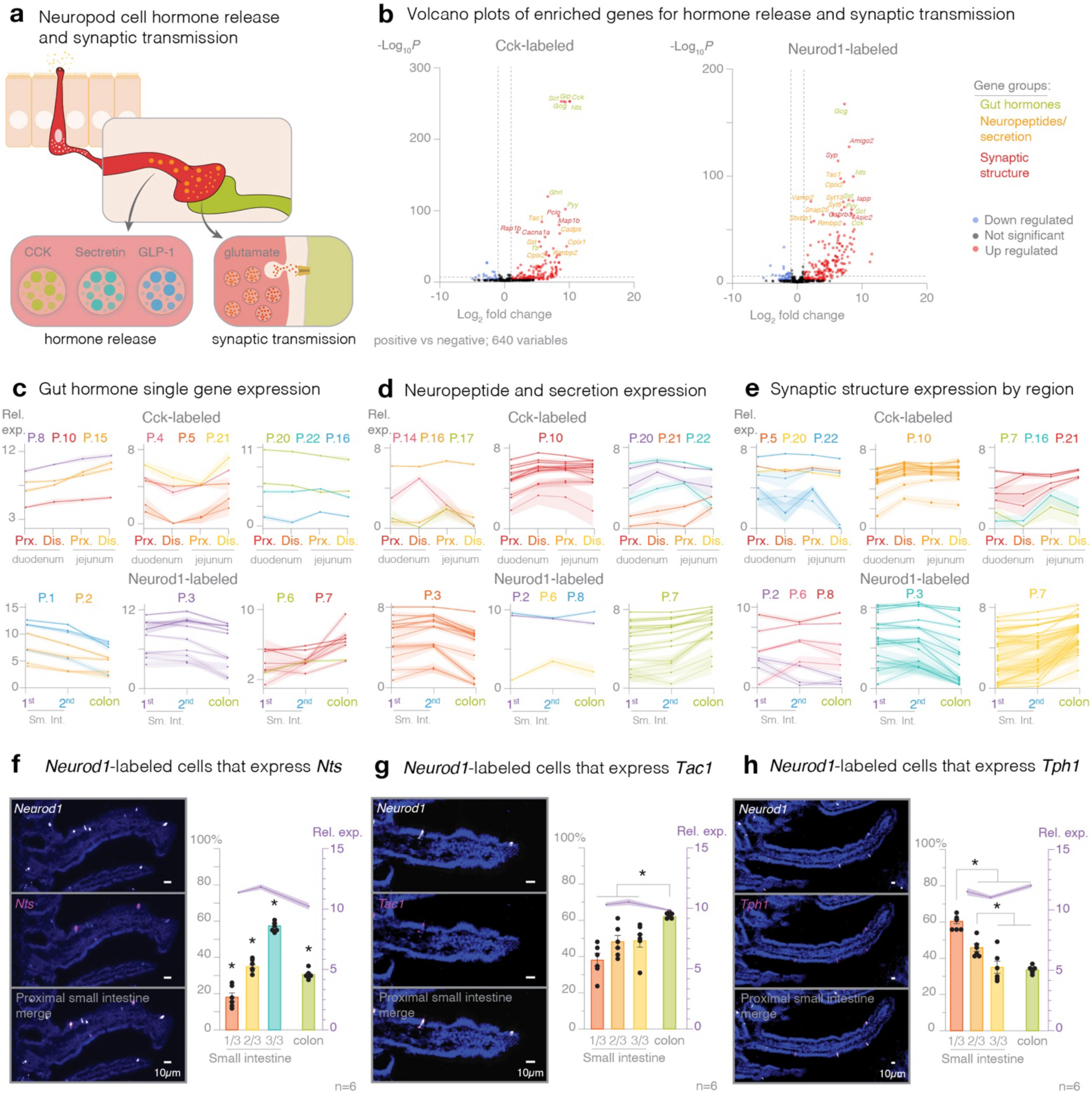
Spatially regulated genes for hormone release, neurotransmitter release, and synaptic stricture. (A) Neuropod cells have dual functionality for releasing hormones, and fast synaptic transmission. (B) Volcano plots of differentially expressed genes from Cck-labeled (*left*) and Neurod1-labeled (*right*) neuropod cells. Individual line plots representing the relative expression and shaded standard error of the mean (SEM) for significant spatially regulated genes associated with (C) gut hormones, (D) neuropeptides, and (E) synaptic structure in (*top*) Cck-labeled and (*bottom*) Neurod1-labeled cells. In situ hybridization (*left*) representative images from the proximal small intestine, (*right*) bars are quantification of overlap with Neurod1-labeled cells in the small intestine (broken into thirds) and the colon; overlay line plot of the corresponding mean expression of Neurod1-labeled cells for neuropeptide genes: (F) *Nts*, (G) *Tac1*, (H) *Tph1* (Mean±SEM; n=6).

Spatial analysis of Cck-labeled cells identified 22 unique spatial expression patterns, with gene counts ranging from 1 to 467 genes (Supp. Fig. 4A). In Neurod1-labeled cells, we identified 8 unique spatial patterns, with a median of 83 genes per pattern (range: 15 to 1,197 genes) (Supp. Fig. 4A). Heatmaps and line plots of spatially regulated genes revealed distinct regional patterns. Hormonal genes *Gip, Cck, Sct, Ghrl*, and *Iapp* decreased expression from proximal to distal, while *Pyy, Sst,* and *Tac1* expression increased distally (Figure 4C, Supp. Fig. 4B,C). In contracts, there were several more neurotransmission and synapse-associated genes that were spatially regulated (Figure 4D-E, Supp. Fig. 4B,C). Generally, these spatial patterns indicate that gut hormones may have broader activity zones, while synaptic transmission is more varied allowing for tighter specialized sensory transduction based on location. To validate the expression patterns and examine single cell resolution, in situ hybridization was performed on small intestine (segregated into thirds) and colon tissue. As expected, *Chgb* was detected in 100% of Neurod1-labeled cells with consistent expression levels across the gastrointestinal tract (Supp. Fig. 4D). The neuropeptide *Nts* cell numbers mirrored its relative expression levels (Figure 4F). However, *Tac1* expression in the colon decreased compared to the small intestine while the number of *Tac1*-expressing cells increased, suggesting that more cells express less transcript in the colon (Figure 4G). Conversely, *Tph1* expression in the colon increased compared to the distal small intestine, while the number of Tph1-expressing cells remained the constant, implying that in the colon the same density of cells expresses more *Tph1* transcript, potentially indicating increased serotonin signaling (Figure 4H). Together these findings suggest a functional division of labor within neuropod cells. Gut hormones may mediate broader, pan regional effects, while the spatially diverse expression of neuropeptides and synaptic structure gene allows for more nuanced and rapid communication, enabling precise regulation of gut sensory transduction.

### Expression of immune-related genes throughout the intestinal tract

Gut sensing is intricately linked to the immune system. The gut is at the intersection of the environment and the internal body, certain foods can promote inflammation, and there is a resident microbe, fungal, and viral community collectively known as the gut microbiome.^11,12^ Volcano plots of genes associated with cytokines, cytokine receptors, and microbiome sensing in both Cck- and Neurod1-labeled cells revealed both up- and down-regulated genes (Figure 5A). Heatmaps of spatially expressed immune-related genes showed fewer spatially regulated genes in gut sensory cells compared to the hormone and neurotransmission groups (Supp. Fig. 5A), suggesting that while immune interactions utilize similar mechanisms throughout the intestine, some regional specificity exists.

**Figure 5.**
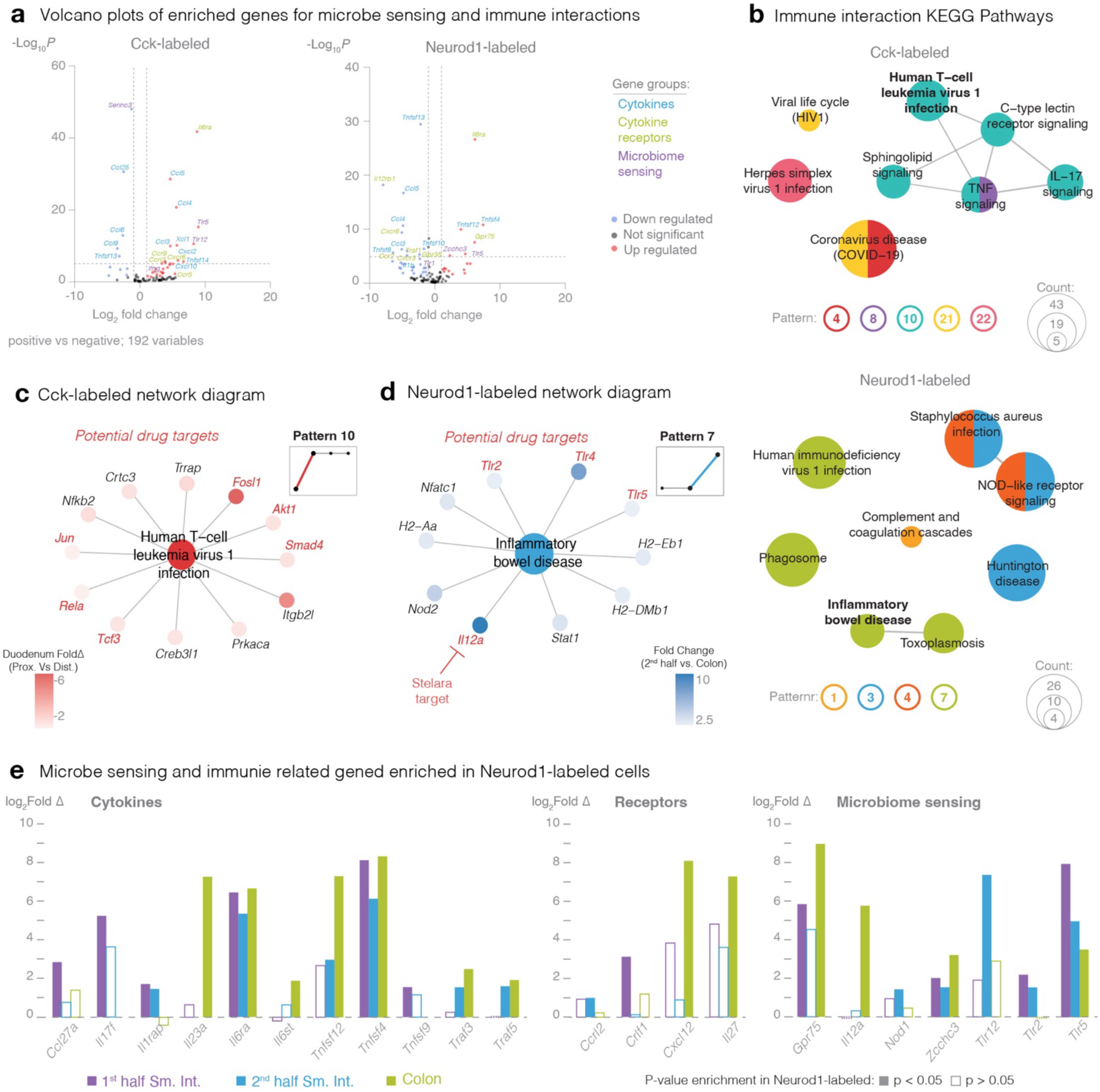
Immune interactions of gut sensory cells throughout the gastrointestinal tract. (A) Volcano plots of differentially expressed genes from Cck-labeled (*left*) and Neurod1-labeled (*right*) neuropod cells. (B) Network enrichment map of KEGG pathways associated with immune interactions colored by spatial patterns of Cck-labeled (*top*) and Neurod1-labeled (*bottom*) cells. (C) Network diagram of genes associated with KEGG pathway Human T-cell leukemia virus 1 infection displayed as fold change of proximal duodenum versus distal duodenum in Cck-labeled cells. (D) Network diagram of genes associated with KEGG pathway Inflammatory Bowel Disease displayed as fold change of 2^nd^ half small intestine versus colon in Neurod1-labeled cells. (E) Log_2_Fold Change of enriched immune interaction genes in Neurod1-positive versus negative cells. Filled bars indicate genes with significant p-value (n≥5).

To investigate regionally specific immune mechanisms, we performed gene over-representation analysis using Gene Ontology (GO) and KEGG pathway analysis. Immune-related GO terms in Cck-labeled cells were primarily associated with increasing Pattern 10, while in Neurod1-labeled cells, they were split between decreasing trend Pattern 3 and increasing trend Pattern 7 (Supp. Fig. 5B). KEGG pathway analysis, while associated with a broader range of patterns, confirmed the trends found in the GO analysis, notably including Cck-labeled Pattern 10 and Neurod1-labeled Pattern 7, both showing increasing trends (Figure 5B). These data highlight a trend of increased immune interaction towards the distal gastrointestinal tract, consistent with the increased density of microbes in this region.^32^

Focusing on genes within highly enriched KEGG terms revealed several potential drug targets under active investigation (Figure 5C,D). One example is *Il12a*, the target of the antagonist drug Stelara, used to treat Crohn’s disease and ulcerative colitis, both inflammatory bowel diseases.^33^ Drug targets within the gut epithelium are particularly attractive due to the potential for localized drug delivery, minimizing systemic exposure and off-target effects.^34^ Furthermore, access to neuropod cells offers the added potential to target brain regions from the gut. Therefore, we focused on immune-related genes specifically enriched and highly expression in neuropod cells (Supp. Fig. 5C). Focusing on the Neurod1-labeled population allowed us to identify targets throughout the entire gastrointestinal tract. Four genes were significantly enriched both the small intestine and colon: *Il6ra, Tnfsf4, Zcchc3,* and *Tlr5* (Figure 5E). While the functional roles of *Il6ra*, *Tnfsf4*, and *Zcchc3* in neuropod cells remain to be elucidated, *Tlr5*, a microbial pattern recognition receptor activated by bacterial flagellin.^35^ The deletion of *Tlr5* in gut epithelial cells leads to weight gain in animals,^36,37^ suggesting that these enriched genes have the potential to influence other gut-related behaviors.

### Nutrient sensing is spatially regulated in neuropod cells throughout the gastrointestinal tract

The main calorie providing nutrient groups are fats, carbohydrates, and protein,^38^ however standard rodent chow is comprised of many essential vitamins and minerals (Figure 6A). These vitamins and minerals also need to be sensed and absorbed, and gut sensory cells are tasked with sensing these diverse nutrients via specific receptors. We analyzed nutrient receptor expression across all nutrient groups. Volcano plots of both Cck- and Neurod1-labeled cells show mixed enrichment for genes associate with fats, sugars, protein, vitamin and mineral sensing (Figure 6B). Heatmaps of spatially expressed nutrient receptor genes revealed that gut sensory cells are tuned to all nutrient groups (Supp. Fig. 6A). Sugar sensing genes were especially enriched in the Cck-labeled cells, suggesting a greater sensory capacity for carbohydrates in the proximal small intestine. Gene Ontology (GO) analysis provided an overview of spatially regulated nutrient-sensing pathways in neuropod cells (Supp. Fig. 6B). In Cck-labeled cells, most GO terms were associated with either the increasing Pattern 10 (highly enriched for carbohydrate/sugar homeostasis), or the decreasing Pattern 20 (enriched for cholesterol homeostasis). This suggests that homeostatic functions are based on early sensing of fatty acids in the proximal duodenum, followed by sugar sensing in the jejunum. In Neurod1-labeled cells, most GO terms were associated with Pattern 3 (high expression in the small intestine, low in the colon), suggesting that general nutrient sensing is concentrated in the small intestine.

**Figure 6.**
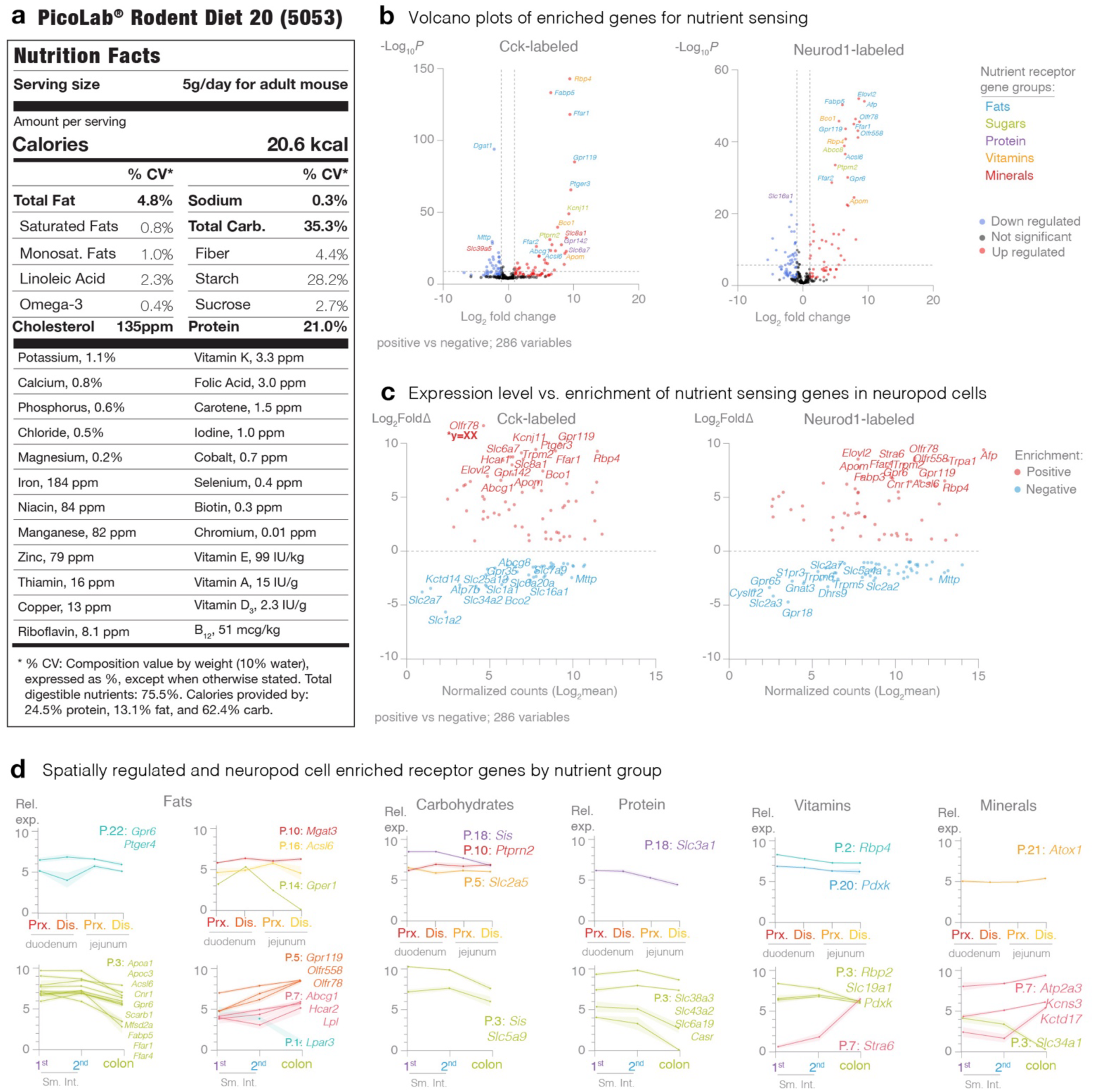
Spatial gene expression based on nutrient groups. (A) Nutrition facts of standard rodent chow. (B) Volcano plots of differentially expressed genes from Cck-labeled (*left*) and Neurod1-labeled (*right*) neuropod cells based on general nutritional groups. (C) Normalized count gene expression levels compared to fold change of differentially expressed nutrient receptor genes. (D) Individual line plots representing the relative expression and shaded standard error of the mean (SEM) for significant spatially regulated genes for nutrient groups, from left to right: fats, carbohydrates/sugars, protein, vitamins, and minerals. Cck-labeled (*top*) and Neurod1-labeled (*bottom*).

As sensory cells, neuropod cells likely play a specialized role in nutrient detection, as opposed to absorption, since they represent only a small percentage of epithelial cells. Therefore, we focused on highly expressed and significantly enriched genes in neuropod cells (Figure 6C). This analysis revealed strong enrichment of fat-sensing genes in both Cck- and Neurod1-labeled cells, indicating a robust role in lipid metabolism. Notably, Cck-labeled cells expressed a higher percentage of sugar receptors compared to Neurod1-labeled cells (Supp. Fig. 6C). In addition to nutrient identity, other food qualities, such as “warm” and “cool,” are detected by transient receptor potential (TRP) channels expressed in gut sensory cells (Supp. Fig. 6D). These sensations, often perceived as “irritants,” have been associated with visceral hypersensitivity.^39^

Next, we examined genes that were both spatially regulated and enriched in neuropod cells to identify key regions for specific nutrient sensing (Figure 6D). Genes associated with sugar transport and metabolism (*Slc2a5, Sis, Slc5a9*) were highly expressed proximally and decreased distally, while the insulin secretion gene *Ptprn2* increased from proximal to distal small intestine, suggesting a shift from sugar sensing to insulin release. Protein sensing genes (*Casr, Slc6a19, Slc38a3, Slc43a2*) tended to be more highly expressed in the middle to distal small intestine, consistent with the region where proteins have been broken down to signal amino acids.^40^ Vitamin sensing was predominately observed in the small intestine, except for Vitamin A sensor gene *Stra6*. Indicating a role for consistent sensing of Vitamin A throughout the gastrointestinal tract. Mineral sensing was mostly confined to the colon, where ion transport is more tightly regulated.^41^ Further research should investigate the functional roles of specific sensory receptors enriched in gut sensory cells and how they influence food choices.

Our findings reveal a spatially organized sensory map of gut sensory cells, where different regions are specialized for detecting distinct signals including immune signals, and nutrients, then responding via hormones, and neurotransmission. This map is analogous to the cortical homunculus in the brain, representing a map of the body’s sensory and motor functions therefore we term it the *intestinal homunculus*. The *intestinal homunculus* reflects the characteristics of our gut sense to differentially encode and transmit information to the brain, influencing various aspects of physiology and behavior. By mapping the transcriptional landscape of gut sensory cells, we have begun to elucidate the complex code by which the gut communicates with the brain. Further research is needed to fully decipher the language of the gut and its implications for human health. In essence, we are just beginning to understand the complex sensory symphony orchestrated by the gut, a symphony that profoundly influences our perception of our resident microbes and the food we eat.

## METHODS

### Animals and Ethics

Mouse care and experiments were carried out in accordance with protocols approved by the Institutional Animal Care and Use Committee at Duke University Medical Center under protocol A280-18-12. Mice were housed in the Duke University animal facilities and kept on a 12-hour light-dark cycle. They received food and water ad libitum. All animals used in this study were 4-6 weeks old. The mouse strains used were: CckCre (Jackson Labs: 012706), Neurod1Cre (Jackson Labs: 028364), and LSL-tdTomato (Jackson Labs: 007914).

### Gut epithelial cell dissociation

The first 12 cm of the small intestine, cut into 3cm segments, of CckCre_tdTomato mice, or the first half small intestine, second half small intestine, and colon of Neurod1Cre_tdTomato mice were collected and epithelial cells were dissociated as previously described.^3^ Briefly, gut was dissected, flushed with PBS, cut open lengthwise and into ∼1cm pieces, before being placed in 400mM EDTA in PBS at 4 °C for 15 minutes, then at 37 °C for 15 minutes. New cold PBS was added, and villi and crypts were mechanically detached and passed through a 100μm filter. Villi and crypts were spun down and further digested in collagenase (400 U/ml) and dispase (50 U/ml). Cells were spun down and re-suspended in L-15 media with 5% FBS and DNase to prevent cell clumping. Cells were passed through a 70 μm then 40 μm cell strainer to isolate single cells. Then, cells were sorted using fluorescence-activated cell sorting (Sony Sorter), selecting for tdTomato fluorescent cells. Equal numbers of tdTomato-negative cells were collected. Cells were sorted into the lysis buffer, and RNA was immediately extracted using the RNeasy Plus Micro Kit (Qiagen) following the manufacturers’ guidelines. Samples were stored at −80°C before library preparation.

### Library preparation, bulk RNA-seq mapping, gene counting, and quality control

Reverse transcription, cDNA pre-amplification, and sequencing library preparation were performed as previously described in [PMID: 35320714]. Briefly, reverse transcription and cDNA pre-amplification were performed using the SMART-Seq v4 Ultra Low Input RNA Kit for Sequencing (Clontech/Takara). cDNA was harvested and quantified with the Bioanalyzer DNA High-Sensitivity kit (Agilent Technologies). Libraries were prepared using the Nextera XT DNA Sample Preparation Kit and the Nextera Index Kit (Illumina) as per manufacturer instructions. Multiplexed libraries were pooled, and paired-end 150-bp sequencing was performed on the Illumina HiSeq 4000 platform at Sidra Medicine [MM1] A standard RNA-Seq pipeline was launched using the bcbio nextgen toolkit (https://doi.org/10.5281/zenodo.5781867). Briefly, sample Fastq files were mapped to the mouse reference genome GRCm38.p6 (GCA_000001635.8) using STAR (version 2.6.1d). Reads were retained only if uniquely mapped to the genome. FeatureCounts (v2.0.0) was used to obtain the number of reads that mapped to each gene. Sample Transcriptome data quality was assessed by evaluating homogeneity and similarity between samples. We used variance stabilizing transformation (vst) and regularized log transformation(rlog) functions (DESeq2 package (v1.26.0) PMID: 25516281) to transform the raw count data, (dist) function to calculate sample-to-sample distances. We plotted heatmaps of the distance matrix along with PCA of 1000 most highly variable genes and the union of the top 1000 most expressed genes to identify putative outlier samples. Of the 52 samples analyzed for the four regions of Cck-labeled and negative, a total of 6 samples were excluded, still maintaining n=5-7 for each sample group. Of the 60 samples analyzed for the three regions of Neurod1-labeled and negative, a total of 17 samples were excluded, still maintaining n=4-9 for each sample group. We also used DESeq2 (v1.26.0) to normalize the raw count matrix according to sequencing depth and RNA composition (median of ratios method).^42^

### Differential Expression Analysis

Sample transcriptome data quality was assessed by evaluating homogeneity and similarity between samples. We used variance stabilizing transformation (vst) and regularized log transformation (rlog) functions (DESeq2 package, v1.26.0) [45] to transform the raw count data and (dist) function to calculate sample-to-sample distances. Differential expression was tested using the DESeq function from the DESeq2 library, which employs the Wald test to determine if the log2 fold change in gene expression across two groups of samples is higher than expected by chance. Finally, DESeq does p-value correction (adjusted p-value) for multiple testing. Genes with a padj<0.05 and |log_2_FC|>1 were considered as differentially expressed.

### Identification of gene spatial expression patterns along the intestine

To analyze gene expression trends along the intestine, we developed a computational pipeline to identify genes with dynamic expression patterns that significantly deviate from a constant level. First, we normalized the data using the Scanpy function *normalize_total* with default parameters. We then applied a filtering step to retain only genes with a nonzero mean expression in at least 6 of the 12 sections. Next, we ranked genes based on their expression variability along the intestinal axis using a model selection approach. Specifically, we compared two polynomial fits: a constant expression model (degree 0) and a quadratic model (degree 2), using Scikit-learn’s PolynomialFeatures(). The difference in Akaike Information Criterion (ΔAIC) between the two fits was computed using the Regscorepy aic() function. For the analysis shown in Figure 1, we selected the genes with a ΔAIC<-10, which yielded 7,179 differentially expressed genes. To test the robustness of our results, we repeated the analysis with a more stringent criterion by selecting the top 5% genes with the highest -ΔAIC; this yielded 1,332 genes, which could be classified in clusters with similar spatial patterns (Supp. Figure 1A, B). For the selected genes, we fitted their expression levels using polynomial regressions with a low strength ridge regularization with scikitlearn make_pipeline() function, choosing the order of polynomial between 2 and 5 and a regularization strength between 0 and 0.9 that minimizes the loss.

To perform clustering, after rescaling all expression levels between 0 and 1, a distance matrix between genes was computed as:

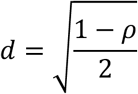

with rho representing the Spearman’s correlation between gene expression trends. The hierarchical clustering linkage matrix was computed using SciPy’s linkage() function with the “Ward” method. Clusters were identified using fcluster(), with the optimal number of clusters estimated via the elbow criterion on the within-sample average silhouette score, computed using Scikit-learn.

### Identification of differentially expressed genes in Neurod1- and Cck-sorted samples

To identify genes with spatially dynamic expression patterns in Neurod1- and Cck-sorted samples, we performed differential expression analysis between adjacent intestinal positions using the following approach. First, we only considered genes detected in at least one spatial location. Differential expression analysis was conducted using pyDESeq2 to compare gene expression between consecutive sections. To select genes with the most significant differential expression, we applied the following thresholds: for Neurod1-sorted samples, we took genes with an adjusted p-value<0.05 and a fold-change greater than 2 or smaller than 0.5; for Cck-sorted samples, the same fold-change threshold was used, p-value was not corrected for multiple testing due to the limited number of samples in the last spatial position. With these criteria, we selected 2270 with samples sorted for Cck, and 2772 for Neurod1-sorted samples. These spatially differentially expressed genes were then grouped based on their expression changes across all pairwise comparisons. This resulted in the identification of 22 groups of genes for Cck-sorted samples and 8 for Neurod1-sorted samples.

### Functional annotation and KEGG pathway

Common genes from each comparison were extracted and used for both Gene over-representation analyses. First, we analyzed GO using the ClusterProfiler package (v4.0.0)^43^. We used Gene Ontology enrichment (enrichGO) and Simplify functions, with ont 1⁄4 “BP”, “MF” or “CC” and pAdjustMethod 1⁄4 “BH” parameters to assess which Biological Processes, Molecular Functions, or Cellular Components were affected in our lists of differentially expressed genes.

The common genes across the different comparisons were annotated by the KEGG database and mapped to KEGG pathways. Based on the total putative identified metabolites as background in each pathway, we used a hypergeometric test to find the significantly enriched KEGG pathways for the differentially accumulated metabolites by R software. The *p*-values were adjusted by FDR with a threshold less than or equal to 0.05.

### Fluorescence in situ hybridization

Wild-type (C57Bl/J6) mice underwent trans-cardiac perfusion with PBS for 3 minutes followed by 4% PFA for 3 minutes at a rate of 600 µl/min. The entire intestine and colon were harvested, opened lengthwise, and divided into different sections: proximal, middle, and distal 1/3 of the small intestine; and colon. Intestinal tissue was then rolled with the proximal end in the center, and post-fixed in 4% PFA for 24 hours. Tissue was then supplemented in 10% sucrose for 1 hour and 30% sucrose for at least 12 hours. Samples were embedded in OCT (VWR) and stored at −80°C. Tissue was sectioned onto slides at 16 μm using a cryostat. RNA detection was performed using the RNAscope Multiplex Fluorescent Reagent Kit v2 Assay (ACD). Briefly, tissue slides were baked for 30 minutes at 60°C, and post-fixed in 10% neutral buffered formalin (VWR) for 60 minutes before being washed in PBS twice (Sigma). Slides were then dehydrated using successive alcohol washes of 50%, 70%, 100%, and a second 100% of ethanol for 5 minutes each. Slides were then incubated with hydrogen peroxide for 10 minutes before undergoing target retrieval using RNAScope reagents in a steamer. Slides were submerged into the RNAScope target retrieval solution at >99°C for 5 minutes. Slides were then treated with Protease III for 30 minutes at 40°C before subsequent hybridization and amplification steps per manufacturer’s instructions. The probes used were all purchased from ACD: Mm-Cck-C2 (cat#402271-C2), Mm-Cck-C3 (cat#402271-C3), Mm-Chgb-C3 (cat#1057371-C3), Mm-Fabp5 (cat#504331), Mm-Neurod1 (cat#416871), Mm-Neurod1-C2 (cat#416871-C2), Mm-Nts-C3 (cat#420441-C3), Mm-Tac1 (cat#410351), Mm-Tph1-O1-C2 (cat#455501-C2), and Mm-Trpv6-C3 (cat#539131-C3). Hybridization signal was detected using Opal dyes (Akoya biosciences) at a dilution of 1:1500, then stained with DAPI (1:4000) for 10 minutes, washed in PBS, and mounted using Fluoro-Gel with Tris Buffer (Electron Microscopy Sciences). Imaging was done on an EVOS Auto 2 (Thermo Fisher Scientific) inverted widefield microscope. Images were adjusted for brightness/contrast using FIJI (Fiji Is Just ImageJ). In each region of the intestine, 100 cells were analyzed from a total of n=6 mice. Cells with >10 puncta within the cell body were considered positive for the gene. Control slides using the negative control probes (ACD) were used to ensure that background staining was <3 puncta per cell. Counts are presented as the mean percentage of co-localization ± S.E.M.

## Data Availability

All data supporting the findings of this study are available within the paper and its Supplementary Information. The mm10 mouse reference genome is available from GENCODE vM23/Ensembl 98.

## Code Availability

All code will be available upon request.

## ACKNOWLEDGEMENTS

The authors would like to thank Drs. Diego V, Bohórquez and Yuanyuan (Kay) He editorial input. Flow Cytometry was performed in the Duke Cancer Institute Flow Cytometry Facility at Duke University, Durham, NC, which is supported by the NCI Cancer Center Support Grant (CCSG) award number P30CA014236. We are thankful to the Integrated Genomics Services of Sidra Medicine for their assistance with RNA-sequencing.

## CONTRIBUTIONS

R.H. and M.M. performed bioinformatics analysis. M.M.K., G.K. and M.A. conducted all animal experiments. M.Z. and A.S. performed spatial pattern analysis. R.H., M. M., L.S., and M.M.K conceptualized and wrote manuscript with input from all authors. L.S. and M.M.K. developed the idea, supervised the project, and acquired funding for the project.

## SUPPLEMENTAL DATA

**Supplemental Figure 1.**
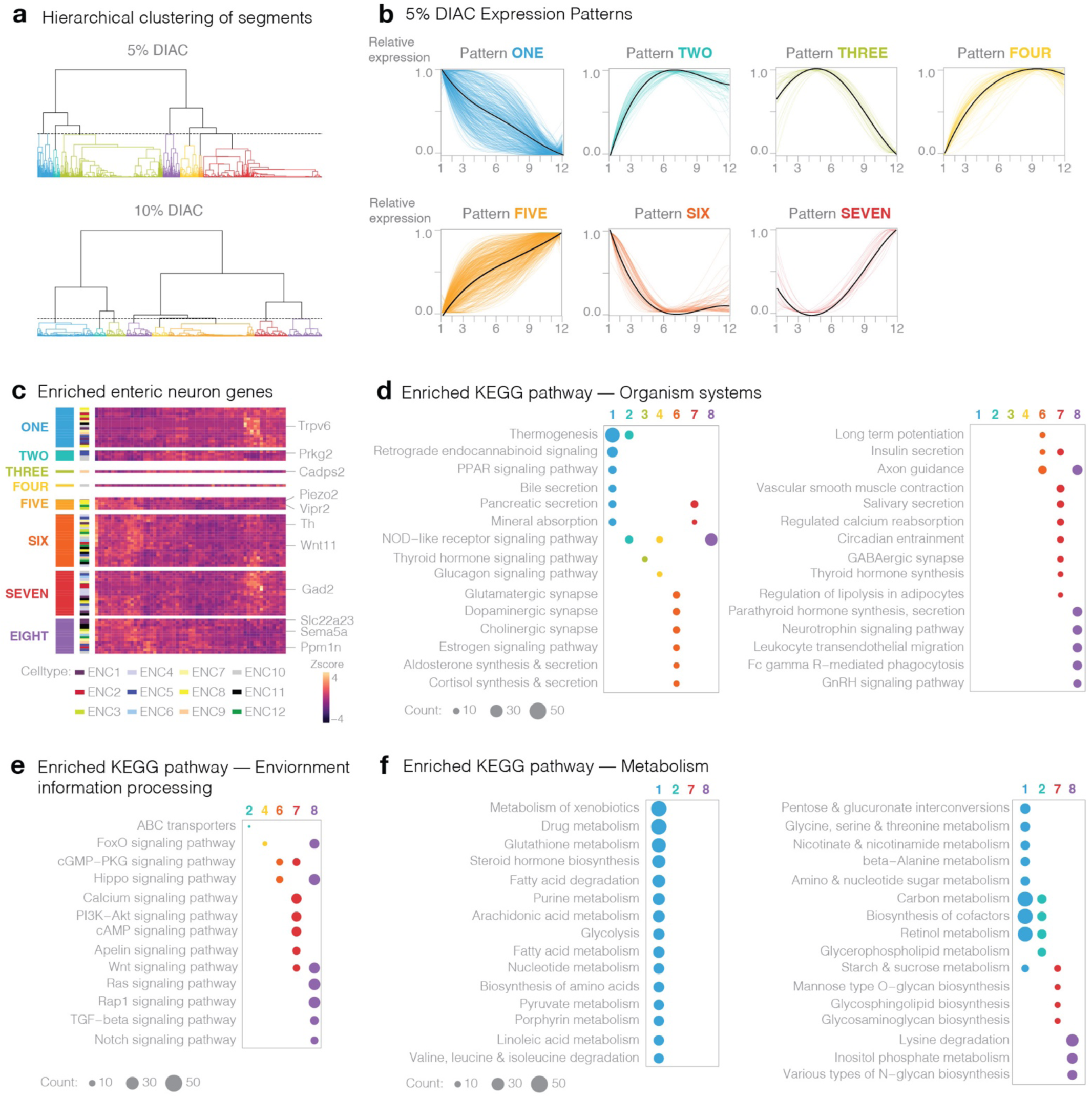
Spatial pattern analysis of small intestinal segments. (A) Dendrogram cuts with optimal cluster numbers, with the resulting regional segment clusters indicated by shaded areas. (B) Spatial transcriptomic analysis using a more stringent 5% cutoff on differential Akaike Information Criteria (-ΔAIC, see Methods), followed by clustering analysis which identified 7 distinct expression patterns. (C) Z-score heatmap of enrichment of Enteric Neuron Class (ENC) genes by expression pattern. KEGG enrichment pathways of: (D) Organismal Systems, (E) Environmental Information Processing, and (F) Metabolism categories. The size of dot represents number of genes enriched, and color indicates an expression pattern.

**Supplemental Figure 2.**
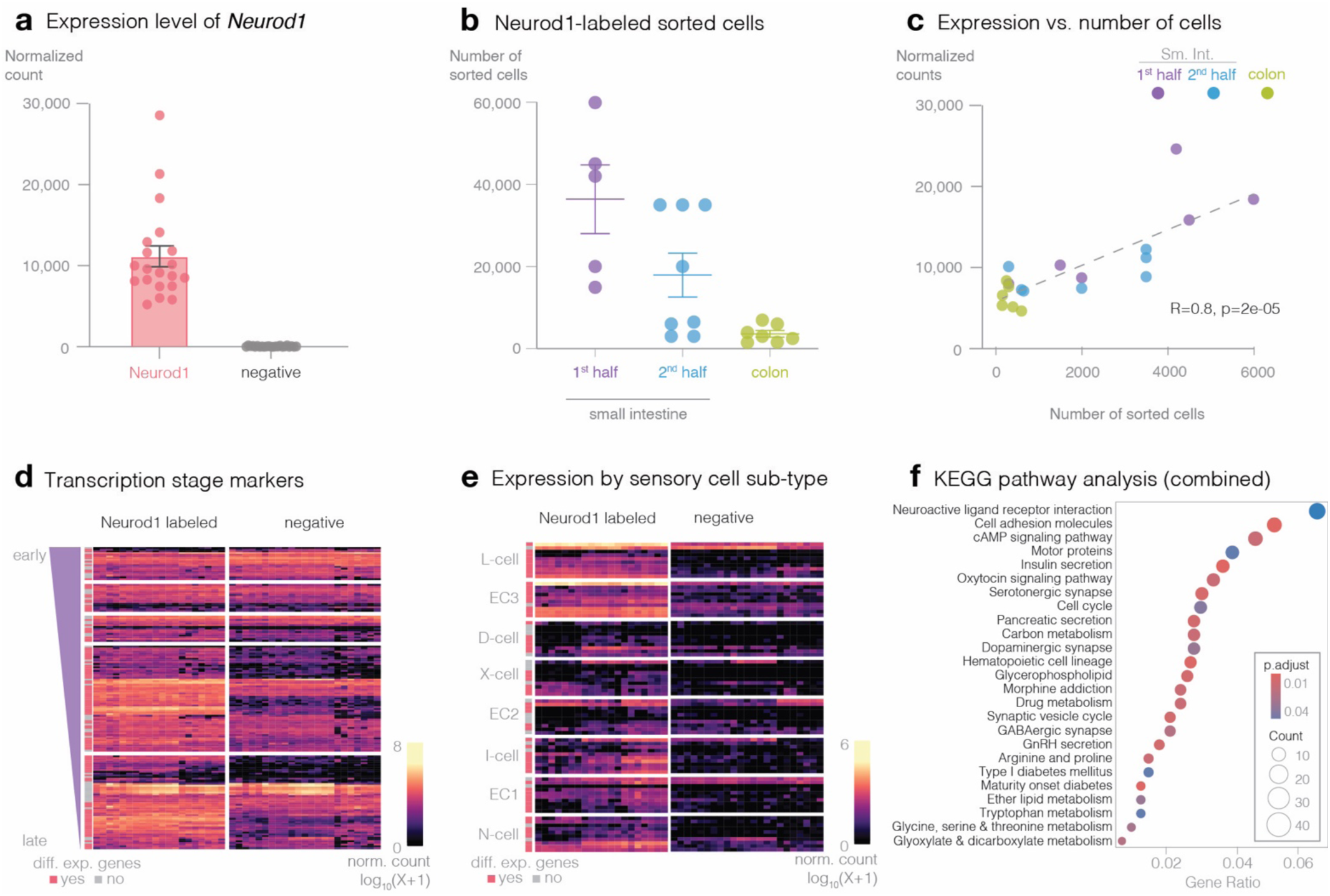
Transcriptional marker of *Neurod1* in sorted cells. (A) Expression level of *Neurod1* in sorted samples (n=4-8 mice per region; mean±S.E.M.). (B) Number of Neurod1-labeled cells sorted by region (mean±SEM). (C) Spearman correlation and linear regression analysis of sorted Neurod1-positive cells and Neurod1 expression level, showing positive correlation (R = 0.8; p = 2e-05). Expression heatmaps of (D) transcriptional stage, and (E) gut sensory cell subtype, in Neurod1-labeled and negative cells across all samples. Color indicates relative expression. (F) Dot plot of enriched KEGG pathways of combined differentially expressed genes within Neurod1-labeled cells (1940 genes). Color indicates significance.

**Supplemental Figure 3.**
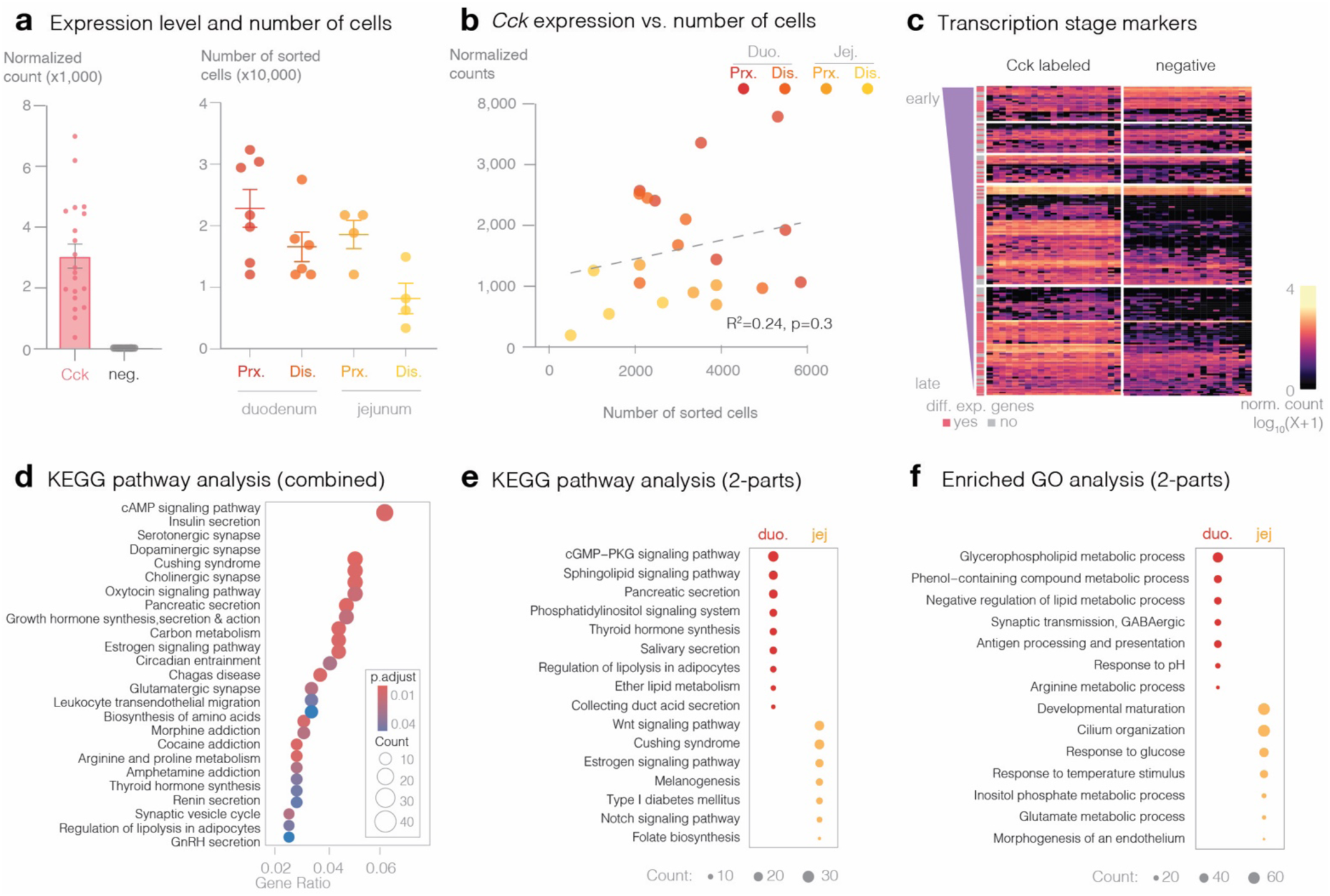
Transcriptional marker of *Cck* in sorted cells. (A) (*left*) Expression level of *Cck* in sorted samples (n=4-7 mice per region; mean±S.E.M.). (*right*) Number of Cck-labeled cells sorted by region. (B) Spearman correlation and linear regression (R2) analysis demonstrating the relationship between the number of sorted Cck-labeled cells and the expression level of *Cck* (R^2^=0.24; p=0.3). (C) Expression heatmap of transcription stage markers in Cck-labeled and negative cells. Color indicates relative expression. (D) Dot plot of top KEGG pathways in combined differentially expressed genes within Neurod1-labeled cells (1940 genes). Color = significance and size = number of genes enriched. Two-part analysis for differentially expressed genes for (E) enriched KEGG pathways, and (F) enriched GO terms. Color = region and size = number of genes enriched.

**Supplemental Figure 4.**
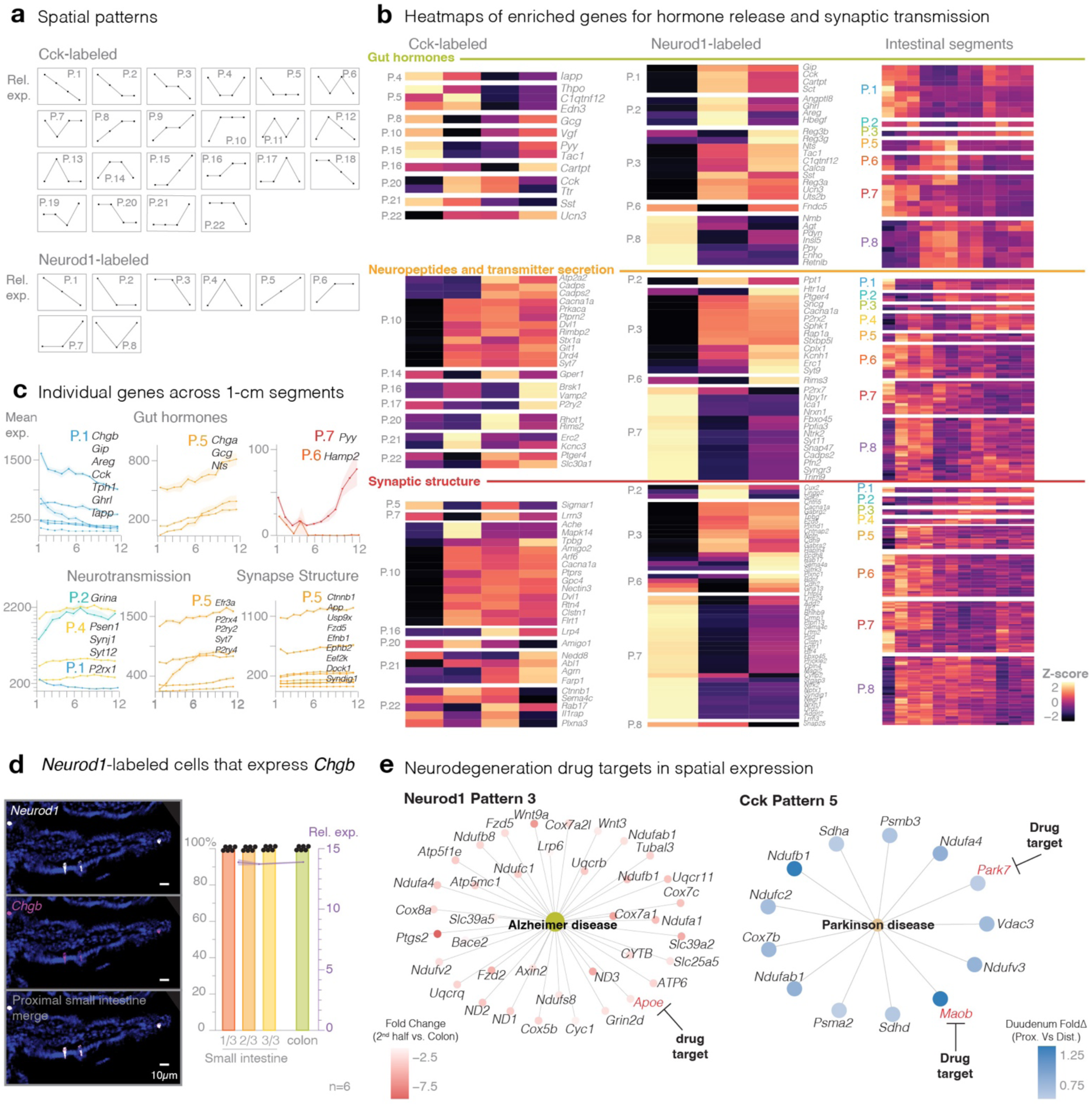
Spatial patterns of neuropod cells through the small intestine and colon. (A) Normalized average expression patterns of spatially differentially expressed genes in Cck-labeled (22 patterns) and Neurod1-labeled (8 patterns) cells along the intestine and colon. The distinct patterns reflect spatial variability in gene expression. (B) Z-score expression heatmap for hormonal, neurotransmission, and synaptic structure genes (from the GOCC database) across Cck-labeled, Neurod1-labeled, and intestinal segment patterns. Clustering is forced based on expression patterns. (C) Individual line plots for gut hormones and neurotransmission genes in 1-cm segments (line=mean relative expression, shading=SEM, n=5). (D) In situ hybridization of *Chgb* (*left*) representative images from the proximal small intestine. (*right*) Bars are quantification of Neurod1-labeled cells that express *Chgb* in the small intestine (broken into thirds) and the colon. Line plot overlay of the corresponding mean expression from the Neurod1-labeled RNA sequencing data. Mean±SEM; n=6. (E) KEGG pathway analysis of disease-related pathways associated with neurodegeneration and drug targets in the context of Neurod1- and Cck-labeled cell patterns. Enrichment results are presented with relevant pathways and candidate targets.

**Supplemental Figure 5.**
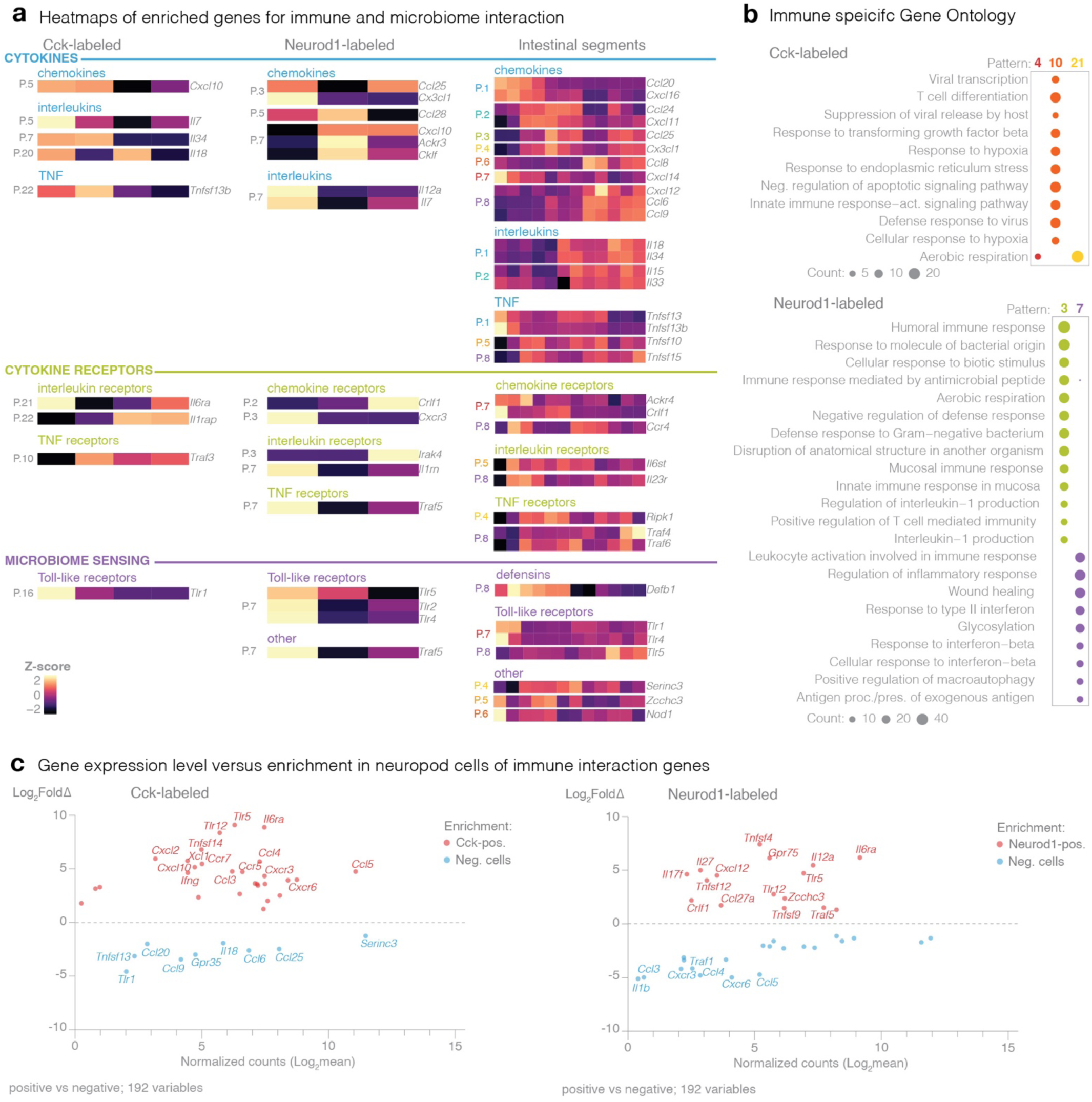
Gene expression levels of enriched genes involved in immune interactions. (A) Expression heatmap Z-score of cytokines, cytokine receptors, and microbiome sensing (GOCC database) genes across Cck-labeled, Neurod1-labeled, and intestinal segments. Clustering based on expression patterns. (B) Immune related gene ontologies in Cck-labeled (*top*) and Neurod1-labeled (*bottom*) cells across their respective spatial patterns (size = number of genes enriched). (C) Normalized count gene expression levels compared to fold change of differentially expressed immune-related genes.

**Supplemental Figure 6.**
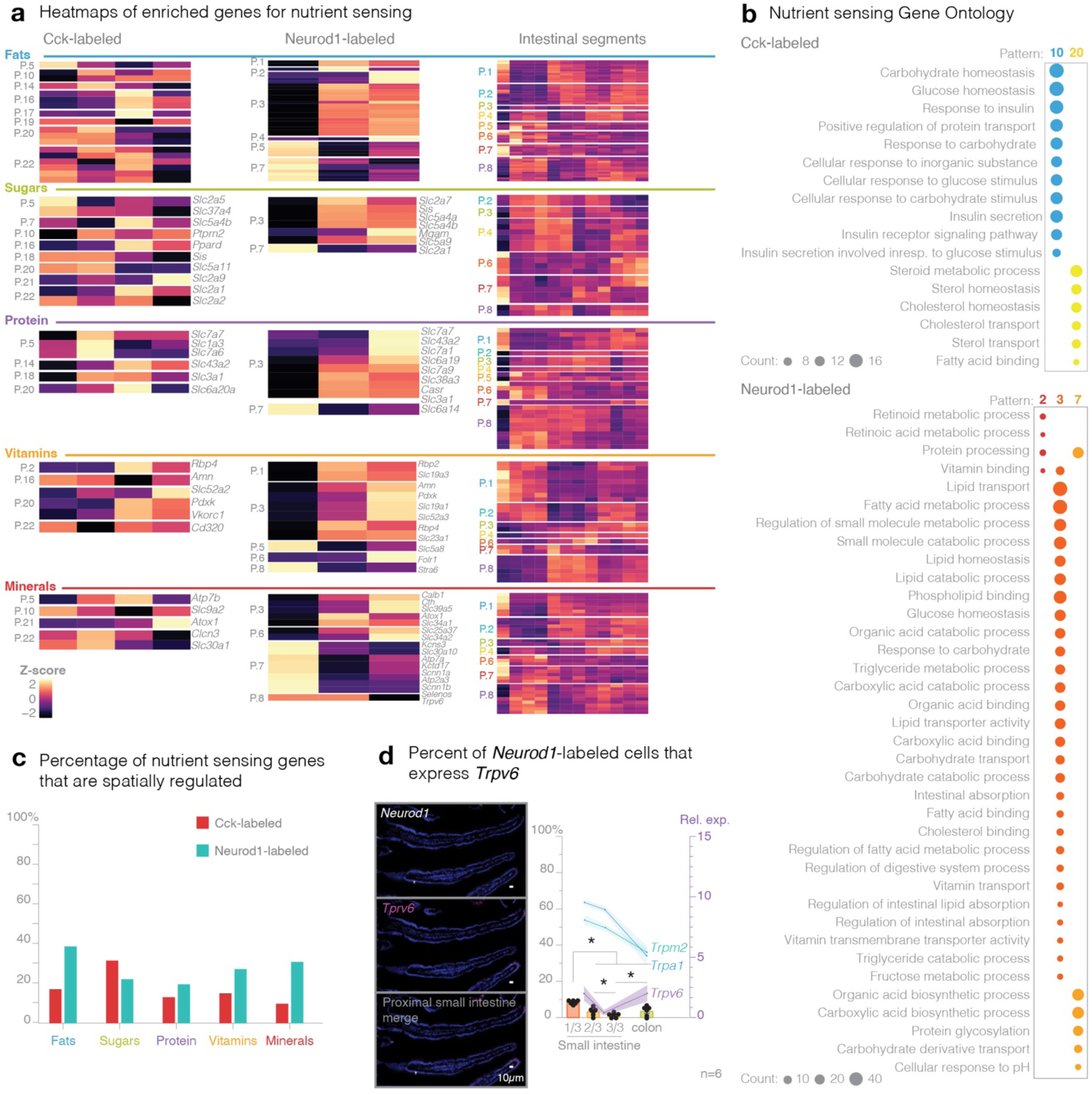
Nutrient sensing gene expression throughout the gastrointestinal tract. (A) Z-score of expression heatmap for genes related to receptors of fats, sugars, proteins, vitamins, and minerals across Cck-labeled, Neurod1-labeled, and intestinal segments. Clustering is forced based on expression patterns. (B) Nutrient sensing related gene ontologies in Cck-labeled (*top*) and Neurod1-labeled (*bottom*) across their respective spatial patterns (size reflects number of enriched genes). (C) Percent of total number of genes per nutrient group expressed in neuropod cells. (D) In situ hybridization (*left*) representative images, (*right*) quantification. Bars are percent overlap with Neurod1-labeled cells in the small intestine (broken into thirds) and the colon; overlay line plot of corresponding mean expression of Neurod1-labeled expression of *Trpm2, Trpa1, and Trpv6* (Mean±SEM; n=6).

## REFERENCES

1. Reimann, F. et al. Glucose sensing in L cells: a primary cell study. Cell Metabol 8, 532–539 (2008).

2. Bellono, N. W. et al. Enterochromaffin Cells Are Gut Chemosensors that Couple to Sensory Neural Pathways. Cell 170, 185–198.e16 (2017).

3. Kaelberer, M. M. et al. A gut-brain neural circuit for nutrient sensory transduction. Science (1979) 361, (2018).

4. Buchanan, K. L. et al. The preference for sugar over sweetener depends on a gut sensor cell. Nat Neurosci 25, 191–200 (2022).

5. Kaelberer, M. M., Rupprecht, L. E., Liu, W. W., Weng, P. & Bohórquez, D. V. Neuropod Cells: Emerging Biology of the Gut-Brain Sensory Transduction. Annu Rev Neurosci 43, 337–353 (2020).

6. Adriaenssens, A. E., Reimann, F. & Gribble, F. M. Distribution and stimulus secretion coupling of enteroendocrine cells along the intestinal tract. Compr Physiol 8, 1603–1638 (2018).

7. Fothergill, L. J. & Furness, J. B. Diversity of enteroendocrine cells investigated at cellular and subcellular levels: the need for a new classification scheme. Histochemistry and Cell Biology vol. 150 693–702 Preprint at 10.1007/s00418-018-1746-x (2018).

8. Gunawardene, A. R., Corfe, B. M. & Staton, C. A. Classification and functions of enteroendocrine cells of the lower gastrointestinal tract. International Journal of Experimental Pathology vol. 92 219–231 Preprint at 10.1111/j.1365-2613.2011.00767.x (2011).

9. Sanchez, J. G., Enriquez, J. R. & Wells, J. M. Enteroendocrine cell differentiation and function in the intestine. Current Opinion in Endocrinology, Diabetes and Obesity vol. 29 169–176 Preprint at 10.1097/MED.0000000000000709 (2022).

10. Kaelberer, M. M. & Bohórquez, D. V. The now and then of gut-brain signaling. Brain Res 1693, 192–196 (2018).

11. Worthington, J. J. The intestinal immunoendocrine axis: Novel cross-talk between enteroendocrine cells and the immune system during infection and inflammatory disease. Biochem Soc Trans 43, 727–733 (2015).

12. Duan, Y. et al. Inflammatory links between high fat diets and diseases. Front Immunol 9, (2018).

13. Worthington, J. J., Reimann, F. & Gribble, F. M. Enteroendocrine cells-sensory sentinels of the intestinal environment and orchestrators of mucosal immunity. Mucosal Immunology vol. 11 Preprint at 10.1038/mi.2017.73 (2018).

14. Strine, M. S., et al. Intestinal tuft cell immune privilege enables norovirus persistence. Sci Immunol 9, (2024).

15. Grün, D. et al. Single-cell messenger RNA sequencing reveals rare intestinal cell types. Nature 525, 251–255 (2015).

16. Hayashi, M. et al. Enteroendocrine cell lineages that differentially control feeding and gut motility. Elife 12, 1–22 (2023).

17. Zwick, R. K. et al. Epithelial zonation along the mouse and human small intestine defines five discrete metabolic domains. Nat Cell Biol 26, 250–262 (2024).

18. Billing, L. J. et al. Single cell transcriptomic profiling of large intestinal enteroendocrine cells in mice – Identification of selective stimuli for insulin-like peptide-5 and glucagon-like peptide-1 co-expressing cells. Mol Metab 29, 158–169 (2019).

19. Gehart, H. et al. Identification of Enteroendocrine Regulators by Real-Time Single-Cell Differentiation Mapping. Cell 176, 1158–1173.e16 (2019).

20. Gerbe, F. et al. Intestinal epithelial tuft cells initiate type 2 mucosal immunity to helminth parasites. Nature 529, 226–230 (2016).

21. Kotas, M. E., O’Leary, C. E. & Locksley, R. M. Tuft Cells: Context- and Tissue-Specific Programming for a Conserved Cell Lineage. Annual Review of Pathology: Mechanisms of Disease vol. 18 Preprint at 10.1146/annurev-pathol-042320-112212 (2023).

22. Silverman, J. B., Vega, P. N., Tyska, M. J. & Lau, K. S. Intestinal Tuft Cells: Morphology, Function, and Implications for Human Health. Annual Review of Physiology vol. 86 Preprint at 10.1146/annurev-physiol-042022-030310 (2024).

23. Morarach, K. et al. Diversification of molecularly defined myenteric neuron classes revealed by single-cell RNA sequencing. Nat Neurosci 24, 34–46 (2021).

24. Li, H. J. et al. Intestinal Neurod1 expression impairs paneth cell differentiation and promotes enteroendocrine lineage specification. Sci Rep 9, (2019).

25. Li, H. J., Ray, S. K., Singh, N. K., Johnston, B. & Leiter, A. B. Basic helix-loop-helix transcription factors and enteroendocrine cell differentiation. Diabetes, Obesity and Metabolism vol. 13 5–12 Preprint at 10.1111/j.1463-1326.2011.01438.x (2011).

26. Hayashi, M. et al. Enteroendocrine cell lineages that differentially control feeding and gut motility. Elife 12, (2023).

27. Gehart, H. et al. Identification of Enteroendocrine Regulators by Real-Time Single-Cell Differentiation Mapping. Cell 176, 1158–1173.e16 (2019).

28. Kato, T. et al. Gene expression of nutrient-sensing molecules in i cells of cck reporter male mice. J Mol Endocrinol 66, 11–22 (2021).

29. Lu, V. B. et al. Adenosine triphosphate is co-secreted with glucagon-like peptide-1 to modulate intestinal enterocytes and afferent neurons. Nat Commun 10, (2019).

30. Haber, A. L. et al. A single-cell survey of the small intestinal epithelium. Nature 551, 333–339 (2017).

31. Fothergill, L. J., Callaghan, B., Hunne, B., Bravo, D. M. & Furness, J. B. Costorage of enteroendocrine hormones evaluated at the cell and subcellular levels in male mice. Endocrinology 158, 2113–2123 (2017).

32. Martinez-Guryn, K., Leone, V. & Chang, E. B. Regional Diversity of the Gastrointestinal Microbiome. Cell Host and Microbe vol. 26 Preprint at 10.1016/j.chom.2019.08.011 (2019).

33. Ahmed, Z. et al. Ustekinumab Induction and Maintenance Therapy in Refractory Crohn’s Disease. New England Journal of Medicine 14, (2019).

34. Vargason, A. M., Anselmo, A. C. & Mitragotri, S. The evolution of commercial drug delivery technologies. Nature Biomedical Engineering vol. 5 Preprint at 10.1038/s41551-021-00698-w (2021).

35. Hu, D. & Reeves, P. R. The Remarkable Dual-Level Diversity of Prokaryotic Flagellins. mSystems 5, (2020).

36. Vijay-Kumar, M. et al. Metabolie syndrome and altered gut microbiota in mice lacking toll-like receptor 5. Science (1979) 328, 228–231 (2010).

37. Vijay-kumar, M. et al. Deletion of TLR5 results in spontaneous colitis in mice. J Clin Invest 117, 3909–3921 (2007).

38. Bayliss, L. C. & Krieger, J. L. Calories in Context: Conceptual Metaphors and Consumers’ Perception and Use of Calorie Information. J Health Commun 23, (2018).

39. Bayrer, J. R. et al. Gut enterochromaffin cells drive visceral pain and anxiety. Nature 616, (2023).

40. Adibi, S. A. & Mercer, D. W. Protein digestion in human intestine as reflected in luminal, mucosal, and plasma amino acid concentrations after meals. Journal of Clinical Investigation 52, (1973).

41. Sellin, J. H. & De Soignie, R. Ion transport in human colon in vitro. Gastroenterology 93, (1987).

42. Ruiz Tejada Segura, M. L., et al. A 3D transcriptomics atlas of the mouse nose sheds light on the anatomical logic of smell. Cell Rep 38, (2022).

43. Wu, T. et al. clusterProfiler 4.0: A universal enrichment tool for interpreting omics data. Innovation 2, (2021).

